# A putative rhamnogalacturonan-II CMP-β-Kdo transferase identified using CRISPR/Cas9 gene edited callus to circumvent embryo lethality

**DOI:** 10.1101/2023.11.15.567147

**Authors:** Yuan Zhang, Deepak Sharma, Yan Liang, Nick Downs, Fleur Dolman, Kristen Thorne, Ian M. Black, Jose Henrique Pereira, Paul Adams, Henrik V. Scheller, Malcolm O’Neill, Breeanna Urbanowicz, Jenny C. Mortimer

## Abstract

Rhamnogalacturonan II (RG-II) is a structurally complex and conserved domain of the pectin present in the primary cell walls of vascular plants. Borate crosslinking of RG-II is required for plants to grow and develop normally. Mutations that alter RG-II structure also affect crosslinking and are lethal or severely impair growth. Thus, few genes involved in RG-II synthesis have been identified. Here we developed a method using CRISPR/Cas9-mediated gene to generate callus carrying loss-of-function mutations in the *MPG2* gene that encodes a putative family GT29 glycosyltransferase. Plants homozygous for this mutation do not survive. We show that in the callus mutant cell walls, RG-II does not crosslink normally because it lacks 3-deoxy-D-manno- octulosonic acid (Kdo) and thus cannot form the α-L-Rha*p*-(1→5)-α-D-kdo*p*-(1→ sidechain. We suggest that MGP2 encodes an inverting CMP-β<u>-K</u>do transferase (RCKT1). Our discovery provides further insight into the role of sidechains in RG-II dimerization.

---

The primary wall surrounding growing plant cells is a dynamic structure comprised primarily of cellulose, hemicellulose and pectin ^1^. The synthesis of these polysaccharides involves a large number of genes, which together may account for up to 10% of a plant’s genome. The Arabidopsis genome encodes at least 567 predicted glycosyltransferases (GTs) across 44 Carbohydrate Active Enzyme (CAZy) families. Many of these GTs are involved in cell wall glycan synthesis. However, only a small number of them have been functionally characterized ^2^.

Rhamnogalacturonan II (RG-II) is a structurally complex and conserved domain of the pectin present in the primary cell walls of all vascular plants. Borate crosslinking of this RG-II is required for these plants to grow and develop normally ^3–7^. Mutations that result in alterations to RG-II structure also affect crosslinking and may be lethal or severely impair growth. Thus, few genes involved in RG-II synthesis have been identified ^8–11^. RG-II has a backbone of homogalacturonan (HG) that is decorated with four structurally distinct side chains (A-D) and two arabinofuranosyl (Ara*f*) substituents (**Fig. 1a**). The side chains are composed of 12 different monosaccharides linked by at least 20 different glycosidic bonds. Thus, RG-II is the most structurally complex polysaccharide yet identified in Nature ^4, 12–14^.

**Fig. 1.**
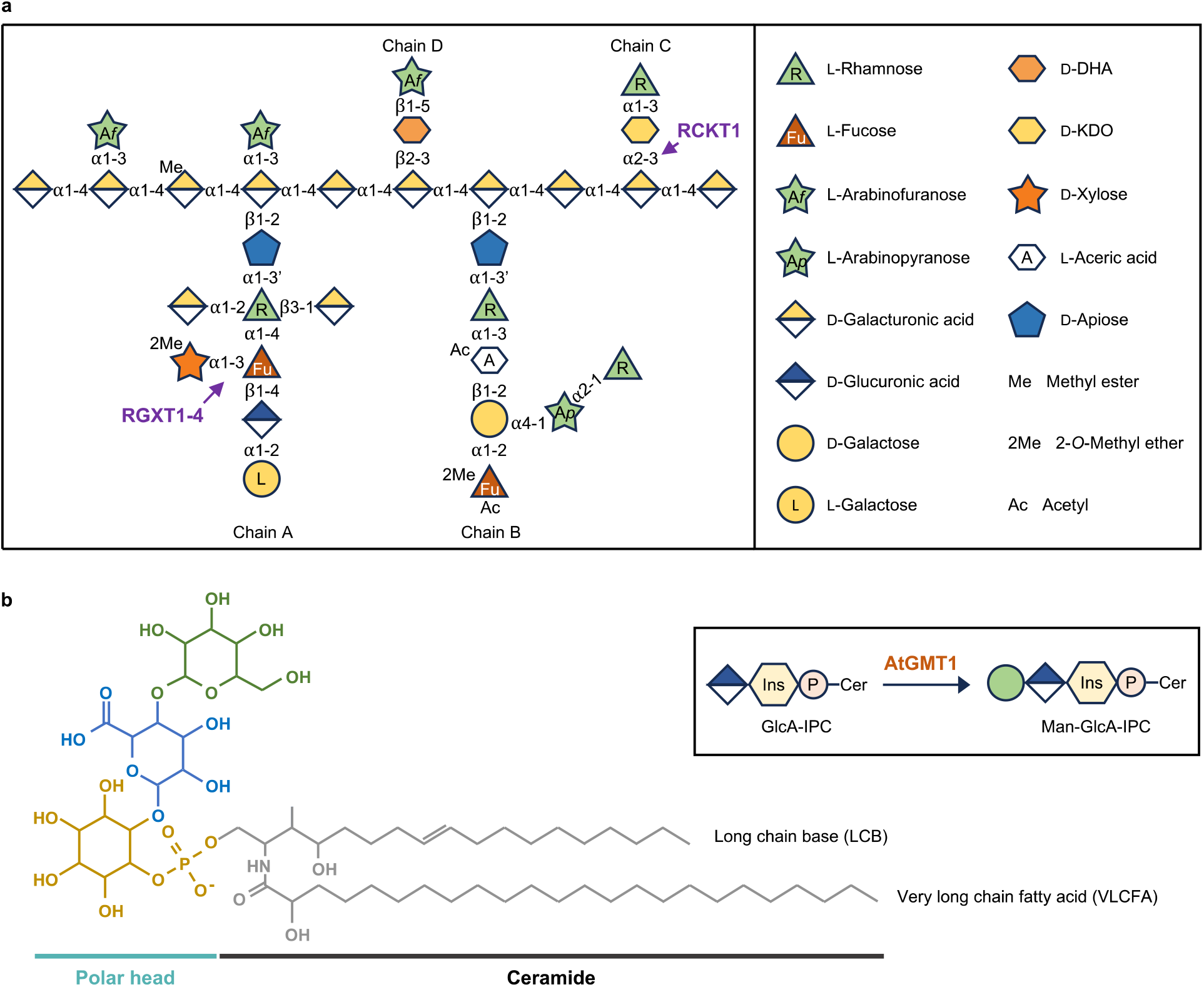
Structure of rhamnogalacturonan-II (RG-II) and Glycosylinositol Phosphorylceramide (GIPC). **a**, Schematic structure of RG-II (modified from Sechet *et al*., 2018). Monosaccharides are shown using the Symbol Nomenclature for Glycans ^84, 85^. **b**, Example structure of a GIPC showing the ceramide (black), inositol phosphate (yellow), glucuronic acid (GlcA, blue), and mannose (Man, green) (modified from Mortimer and Scheller, 2022). Man-GlcA-IPC is the dominant headgroup structure in wild-type Arabidopsis callus tissue. Inset: The synthesis of Man-GlcA-IPC catalyzed by Arabidopsis GMT1 (AtGMT1). Man and GlcA are shown using the same Symbol Nomenclature for Glycans as in **a**.

Plants carrying mutations in genes that affect RG-II synthesis and structure typically exhibit severe growth defects. For example, the Arabidopsis *mur1* mutant, which is impaired in its ability to synthesize GDP-L-fucose (Fuc), is unable to form a normal side-chain A and has a dwarf phenotype ^15, 16^. Similar phenotypes were obtained by silencing the major Golgi-localized GDP-L-Fuc transporter in the *gfp1* mutant ^17^, and by mutations affecting the synthesis and transport of other monosaccharides required to form RG-II, including GDP-L-galactose (Gal) ^18, 19^; UDP-D-glucuronic acid (GlcA), a precursor for UDP-D-apiose (Api) ^20^; UDP-D-Api/UDP-D-xylose (Xyl) ^21^; and CMP-β-3-deoxy-D-manno-octulosonic acid (Kdo) ^22^.

Identifying the GTs involved in the formation of RG-II is required to understand the role of this polysaccharide in cell wall integrity and plant growth. Reverse genetic approaches have had limited success in identifying RG-II-specific GTs. This is primarily due to the deleterious effects that even minor modifications to RG-II structure have on plant growth. To date only two GTs have been implicated in RG-II synthesis. Rhamnogalacturonan Xylosyltransferase (RGXT) 1-4, a family of GTs is believed to add xylose to side chain A ^23–26^. Galacturonosyltransferase 4 (GAUT4) may be responsible for forming the homogalacturonan backbone ^27^. Loss-of-function mutations in GAUT4 and RGXT4 are lethal. Thus, alternate strategies were used to gain insight into their functions: RNA interference (RNAi) knockdown of *GAUT4* in switchgrass and poplar caused a reduction in the amounts and crosslinking of RG-II ^27^; A pollen-rescue strategy enabled the generation of a homozygous RGXT4 (also known as MGP4) loss-of-function mutant, which showed decreased xylose content and reduced RG-II dimerization ^26^. Nevertheless, both strategies have their limitations. Knockdown of target gene expression may be insufficient and obscure the consequences of gene loss-of-function. The pollen rescue strategy is effective only if lethality is associated with the defective male gametophyte. Both methods require extensive crossbreeding and screening yet offer no assurance of achieving stable lines. Hence, there is a need to establish a facile system to generate viable plant lines lacking expression of functional GTs.

Glycosylinositol phosphorylceramides (GIPCs) are an important class of plasma membrane lipids in plants (**Fig. 1b**) ^28^. Previously, we observed that plants carrying mutations that affect the glycosylation of GIPCs had severe growth defects, yet their null mutant callus grew well ^29–31^ . GIPCs have been proposed to have a role in cell-cell adhesion ^31, 32^ as has the formation of a pectin network by borate crosslinking of RG-II. For example, reduced RG-II crosslinking in the *mur1* mutant results in defective cell adhesion, specifically at the interface between stele and cortex cells ^33^. The “*rotten ear*” maize mutant also exhibits decreased cell adhesion as well as disrupted cell wall organization in the ear that has been attributed to incomplete borate cross- linking of RG-II ^34^. This led us to hypothesize that engineered callus tissue may provide an opportunity to identify genes responsible for glycan synthesis that are harmful to normal plant growth and development. Thus, we developed CRISPR-mediated tissue transformation to generate callus carrying loss-of-function GT mutations. We validated this approach with the well characterized GIPC mannosyl-transferase1 (AtGMT1) (**Fig. 1b**) ^30^. We then generated a loss-of-function mutant of a CAZy family GT29 protein (Male gametophyte defective 2, MGP2) currently annotated as a sialyltransferase-like protein in TAIR (arabidopsis.org). The callus mutant together with NMR spectroscopic and mass spectrometric analysis of its RG-II provided evidence that MGP2 is the CMP-β-Kdo transferase (RCKT1) required to form the α-L-Rha*p*-(1→5)-α-D-Kdo*p*- (1→ side chain of RG-II.

## Results

### Arabidopsis GT29 family members are highly co-expressed with primary cell wall- and GIPC-related genes

To identify GTs implicated in cell-to-cell adhesion processes, we used bioinformatics and publicly available arrays of 113 developmental RNA seq datasets, representing 11 distinct tissue and organ types to identify genes that are co-expressed with Arabidopsis *GMT1* ^35, 36^. Disrupting this gene causes severe plant growth defects and reduces cell-cell adhesion ^30, 32^. We then used SUBA (Subcellular Location Database for Arabidopsis Proteins) to refine our list to 75 candidates with likely Golgi localization (**Supplementary Data 1**). These candidates were regarded as putative GTs relevant to cell-cell adhesion. Somewhat unexpectedly, all three Arabidopsis family GT29 members - At1g08280 (GALT29A), At1g08660 (RCKT1, also known as MGP2), and At3g48820 (SIA2) - were included. They are annotated as sialyltransferase-like (ST-like) proteins since they possess motifs characteristic of mammalian family GT29 STs. However, the plant enzymes have no discernible ST activity ^37^. This is not entirely unexpected since plants produce little if any sialic acid ^38^.

We then considered the possibility that the plant GT29 enzymes use a nucleotide sugar donor substrate structurally related to CMP-Sialic acid, the donor for STs. Indeed, *RCKT1* and *SIA2* have been proposed to encode enzymes that catalyze the transfer of 2-keto-3-deoxy-D-manno-octulosonic acid (Kdo) or 2-keto-3-deoxy-D-lyxo-heptulosaric acid (Dha) ^39, 40^. The activated form of Dha remains to be identified. CMP-β-Kdo is the activated form of KDO and the family GT29 proteins utilize an inverting mechanism for glycosyl transfer, indicating they would be able to form the α-2,3-bond that links Kdo to the RG-II backbone and initiates formation of sidechain C (**Figure 1**).

### Creating biallelic mutant calli carrying GT knockouts

No homozygous mutants of *GALT29A*, *RCKT1*, (*MGP2*), and *SIA2* have been isolated, suggesting they are lethal or severely harm the mutant plants. This also suggests that these GTs have an essential role in plant development. This led us to develop a CRISPR/Cas9-mediated tissue culture transformation strategy to generate loss-of-function callus to study the function of GTs. CRISPR/Cas9 technology is a powerful tool for targeted genome editing in plants and has considerable potential for generating knockout mutants. Here, we optimized a tissue culture-based transformation method ^41^, and used it to introduce the CRISPR/Cas9 system with the aim of circumventing the need to produce whole plants and viable seeds (**Supplementary** Fig. 1). The isolated clonal knockout lines can be indefinitely propagated as callus tissue, which provides material suitable for biochemical and functional characterization of plants with primary cell wall chemotypes.

We used a modular cloning system ^42^, which includes a T-DNA destination vector, a Cas9 entry vector, and a guide RNA (gRNA) entry vector (**Supplementary** Fig. 2). To increase the rate of editing, each construct contained three sgRNAs for the target gene (**Fig. 2a, Supplementary Table 1**). Since CRISPR/Cas9 editing typically generates small insertions and deletions (indels) through non-homologous end joining (NHEJ), the disruption of gene function primarily stems from frameshift mutations. To maximize the impact of such disruptions, the sgRNAs were designed to target the protein-coding region proximal to the 5’-untranslated region (5’-UTR).

**Fig. 2.**
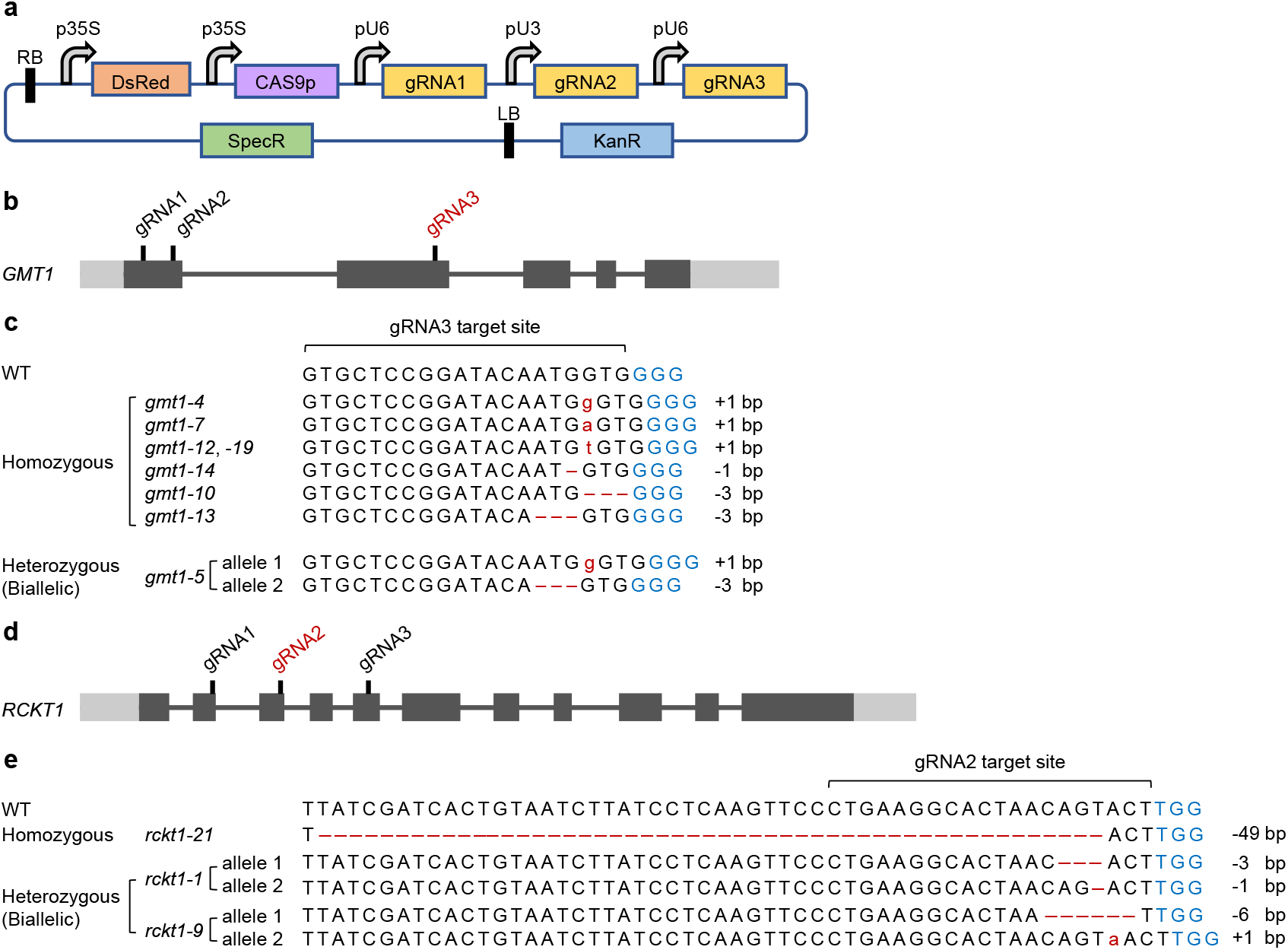
Creation of CRISPR/Cas9-edited Arabidopsis callus lines with different mutations on *GMT1* and *RCKT1*. **a**, Illustration of the T-DNA transformation vector used for CRISPR editing of the target gene. Arabidopsis U6 (AtU6) promoter is used to express gRNA1 and gRNA3, while Arabidopsis U3 (AtU3) promoter is used to express gRNA2. **b**, Schematic diagram of the *GMT1* gene structure with gRNA targeting sites labeled. The black boxes represent exons; the grey boxes represent 5’-UTR and 3’-UTR; the black lines represent introns. CRISPR/Cas9 editing was only detected at the gRNA3 (red) target site. **c**, Gene-editing events at the gRNA3 target site of the *gmt1* callus mutants. Seven homozygous lines were identified: five are *gmt1* knockout mutants with frameshifted and truncated GMT1 protein, and two are mutants with the loss of a single amino acid from the protein sequence. One heterozygous/biallelic line was identified with two mutant alleles, one with a 1 bp insertion causing premature termination of the GMT1 protein, and the other with a 3 bp deletion resulting in the loss of one amino acid from the protein sequence. **d**, Schematic diagram of the *RCKT1* gene structure with gRNA targeting sites labeled. CRISPR/Cas9 editing was only detected at the gRNA2 (red) target site. **e**, Gene-editing events at the gRNA2 target site of the *rckt1* callus mutants. The homozygous knockout mutant *rckt1-21* has a 49 bp deletion within the *RCKT1* gene, creating a frameshifted and truncated RCKT1 protein. The *rckt1-1* heterozygous/biallelic line has two mutant alleles, one with a 3 bp deletion causing the loss of one amino acid from the RCKT1 protein, and the other with a 1 bp deletion resulting in frameshift and premature stop codon in the protein sequence. The *rckt1-9* heterozygous line has two mutant alleles, one with a 6 bp deletion causing the loss of two amino acids from the RCKT1 protein, and the other with a 1 bp insertion resulting in frameshift and premature stop codon in the protein sequence.

### Validating the system with a known GT as the target for knockout

We first targeted *AtGMT1* to test the effectiveness of our CRISPR/Cas9 toolbox and tissue culture transformation method in creating GT knockout mutant calli. W*e* targeted three unique sites in *AtGMT1*: two in Exon1 and one in Exon2 (**Fig. 2b**). The primary roots of ten-day-old Arabidopsis seedlings were used for *Agrobacterium*-mediated transformation, yielding 16 viable calli out of 300 infected root explants (5.3%). The stable integration of the CRISPR/Cas9 cassette in these transformants was indicated by their resistance to kanamycin and the detection of DsRed fluorescence, and further validated via PCR.

To characterize the editing events in each transformant, the genomic region spanning the three target sites was amplified by PCR, Sanger sequenced, and then analyzed using DECODR v3.0 (https://decodr.org/) ^43^. Cas9-induced mutations were introduced into 13 lines (81%) at the sgRNA3 target site in Exon2. This gave us 7 homozygous mutants (with two identical mutant alleles), 1 biallelic mutant (with two distinct mutant alleles) and 5 chimeric mutants (carrying two or more types of alleles, with or without mutation) (**Fig. 2c**). The 7 editing events detected in the homozygous lines consisted of 5 frameshifting indel mutations, expected to result in truncated proteins, and 2 in-frame deletion mutations that would lead to the loss of one amino acid (**Fig. 2c**). The single biallelic mutant had one allele with a frameshift insertion, and another with a frame-preserving deletion (**Fig. 2c**). All the transgenic lines were unedited at the sgRNA1 and sgRNA2 target sites of Exon1, while 3 lines completely escaped targeted editing of *AtGMT1* by the CRISPR/Cas9 complex. Thus, our strategy enables the efficient creation of clonal knockout lines in the form of callus, where every individual cell carries the same frameshift mutation within the target gene.

### Validating the function of *gmt1* by GIPC analysis

We analyzed the GIPCs of the callus lines to assess the suitability of homozygous frameshift mutant calli for decreasing or eliminating GT activity. A transgenic callus line without CRISPR editing was included as a non-edited (NE) control. We also examined callus derived from Col-0 seedling roots and a *gmt1-3* T-DNA insertional mutant.

GIPCs were extracted from callus tissue grown in liquid media and the degree of glycosylation was determined using thin layer chromatography (TLC) ^44^. The GIPC profiles of the *gmt1-4* and *gmt1-3* mutants were clearly different from those of Col-0 and the NE control (**Supplementary** Fig. 3). Notably, the GIPC bands exhibited a larger shift in both *gmt1* mutants, indicating a reduction in molecular weight due to the absence of mannosylation ^30^. Furthermore, the bands displayed a compact and tightly clustered appearance in contrast to the dispersed bands observed in the negative control lines. This distinctive feature marked a significant decrease in the diversity of GIPC species when normal mannosylation of the sugar head group is absent.

We next determined the mannose content of the extracted GIPCs. The monosaccharides were released by hydrolysis with trifluoroacetic acid (TFA) and quantified by high-performance anion exchange chromatography coupled with pulsed amperometric detection (HPAEC-PAD) ^30^. The mannose content was substantially reduced (3-7 fold) in both *gmt1* mutants compared to that in the wild-type and NE control (**Supplementary Table 2**). Our qualitative TLC assay and glycosyl residue composition analysis demonstrate that the *gmt1* CRISPR callus mutant recapitulates the GPIC chemotype of the *gmt1-3* T-DNA insertional mutant.

### Creating homozygous *RCKT1* knockout mutant callus

We next targeted the *RCKT1* gene identified in our bioinformatic screen. The tri-gRNA module in the transformation vector was designed to target 3 distinct sites within the *RCKT1* coding region (Exon 2, 3, and 5) (**Fig. 2d**). We obtained 16 viable calli out of 410 infected root explants (3.9% efficiency). Cas9-mediated editing was restricted to the sgRNA2 target site of *RCKT1*. Genotyping of the transgenic lines identified the presence of 1 homozygous mutant, 2 heterozygous/biallelic mutants, 11 chimeric mutants, and 1 NE control line. The homozygous mutant *rckt1-21* carried a “-49bp/-49bp” biallelic mutation, disrupting the open reading frame and introducing an early stop codon (**Fig. 2e**). Consequently, the predicted product of the *RCKT1* mutant allele in *rckt1-21* is a truncated protein consisting of 104 amino acid residues at the N-terminus, with 73 residues preserved and 31 altered. This mutant RCKT1 variant is expected to lack the catalytic domain, rendering *rckt1-21* an ideal knockout line for cell wall analysis. In contrast, the two heterozygous mutants, *rckt1-1* and *rckt1-9*, harbored a “-1/-3” and “+1/-6” biallelic mutation, respectively (**Fig. 2e**). These mutations resulted in one allele carrying a frameshift indel mutation (−1 or +1), while the other allele containing a frame-preserving deletion (−3 or −6), which led to the loss of one or two amino acid(s). It’s unknown whether such biallelic mutations would cause the complete loss of *RCKT1* function in these heterozygous mutants. The chimeric mutants possessed wild-type alleles at varying percentage (6-52%), which remained recalcitrant to CIRSPR/Cas9 editing.

The homozygous *rckt1-21*, the heterozygous/biallelic *rckt1-1* and *rckt1-9* are gene edited lines that can be used to study AtRCKT1 function. The non-edited *rckt1-6*, with a T-DNA insertion but lacking any *AtRCKT1* editing is a control.

### The absence of RG-II Kdo from the *rckt1* mutant

To explore the effect of *RCKT1* mutation on cell wall glycosyl residue composition, we grew the *rckt1* callus lines and controls in liquid media, and then isolated cell walls for all lines as alcohol-insoluble residue (AIR). The monosaccharide compositions of the walls from the *rckt1* mutants and the wild-type and the non-edited control lines were similar (**Fig. 3a-b, Supplementary Data 2**). However, we were unable to detect Kdo in intact AIR, thus we isolated RG-II from the callus walls. To this end we treated the AIR with an endopolygalacturonase (EPG) and separated the solubilized products by size-exclusion chromatography (SEC) ^45^ to obtain an RG-II rich fraction (**Fig. 3c**). This procedure also shows the relative abundance of the RG-II monomer and dimer present in the walls at the time of extraction. The dimer accounts for ∼80% of the RG-II in Col-0 and the NE control (**Fig. 3c**, **Supplementary Table 3),** a result consistent with previous reports ^9, 19, 46^. However, in both the homozygous *rckt1-21* mutant and the heterozygous/biallelic *rckt1-9* mutant the monomer accounted for at least 74% of the RG-II (**Fig. 3c, Supplementary Table 3**), showing that the ability to form the dimer is diminished in the *rckt1* mutants.

**Fig. 3.**
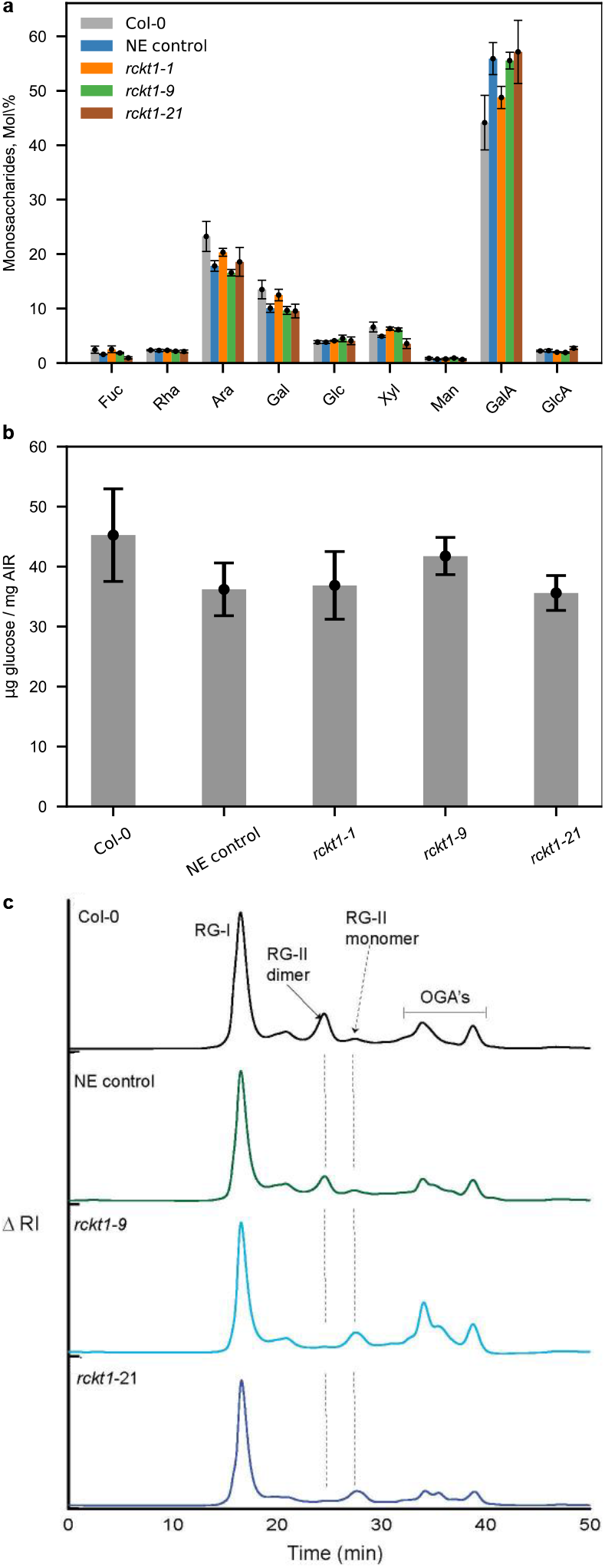
Cell wall polysaccharides and pectin analysis of *rckt1* callus mutants. **a**, Monosaccharide composition of non-cellulosic cell wall polysaccharides **b**, Cellulose-derived glucose. Data in **a** and **b** are means of three biological replicates ± SD. No statistically significant difference (ANOVA test) was observed between Col-0 wild-type, Non-edited (NE) Control, and the homozygous and heterozygous/biallelic *rckt1* mutants. **c**, Separation of pectic domains using size-exclusion chromatography (SEC). The representative SEC profile of endopolygalacturonase (EPG) and pectin methyl esterase (PME) treated alcohol-insoluble residues (AIR) are shown for Col-0 wild-type (black), NE Control (green), *rckt1-9* (light blue), and *rckt1-21* (dark blue). The position of rhamnogalacturonan I (RG-I), RG-II dimer and monomer, and the oligogalacturonides (OGAs) are shown.

The absence of Kdo in the RG-II from the two *rckt1* mutants was first established by solution state nuclear magnetic resonance spectroscopy (NMR). To facilitate the detection of signals corresponding to Kdo and Dha, the RG-II (dimer) was treated with dilute HCl to generate the monomer. Resonances diagnostic for Kdo (H3, δ 1.88 ppm) and Dha (H3, δ 1.78 ppm) were clearly discernible in the wild-type RG-II (**Fig. 4a-b**). The Kdo signal was absent in the *rckt1* mutants (**Fig. 4c-d**). A 2D TOCSY NMR experiment showed that only the wild-type RG-II contained resonances characteristic of H3 and H3’ (δ 1.88 and 2.13 ppm) correlated with H4 (δ 4.13) of Kdo, whereas the Dha H3 and H3’ resonance (δ 1.78 and 2.05 ppm) correlated with H4 (δ 4.08) were present in both the controls and the *rckt1* mutants (**Fig. 5**).

**Fig 4.**
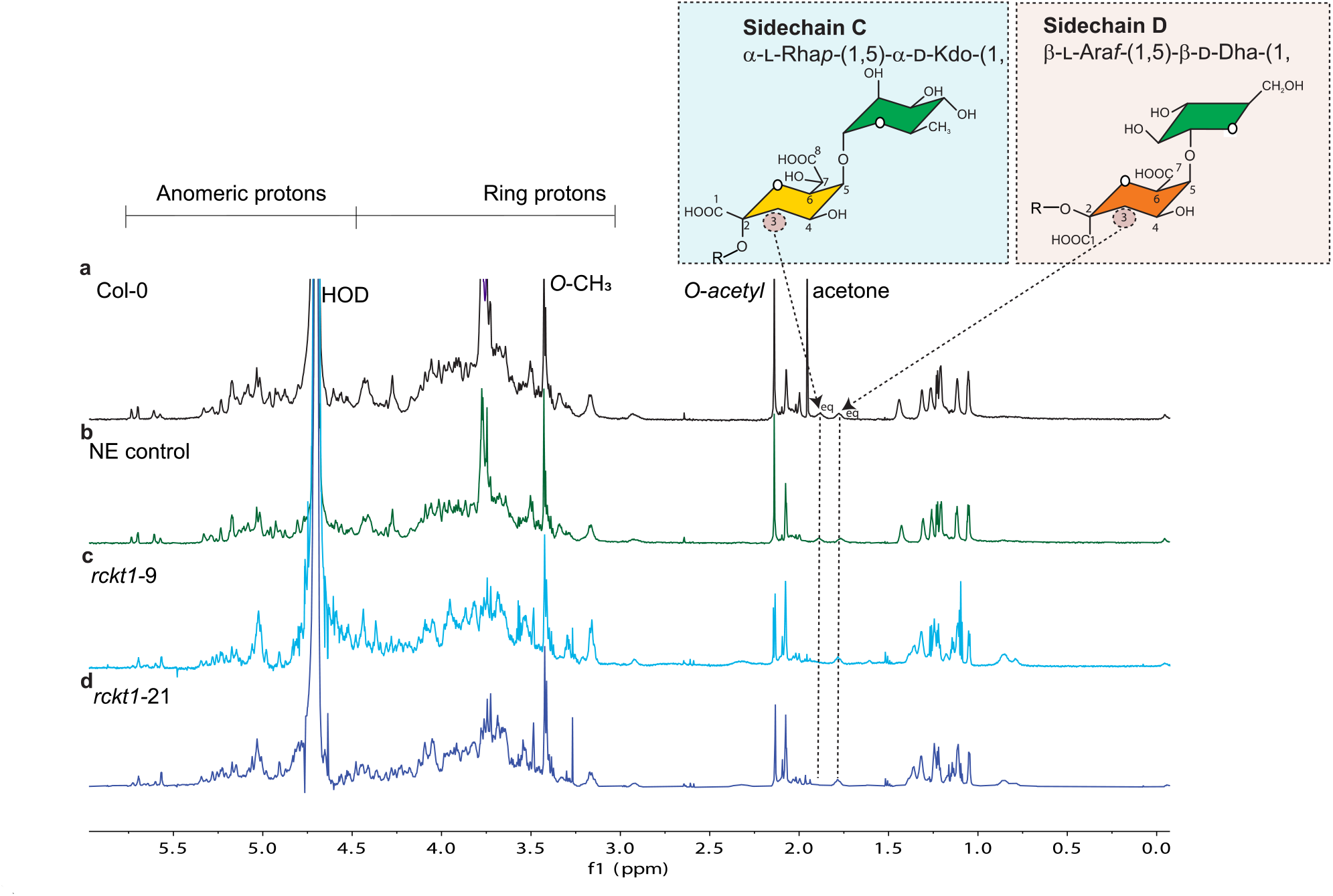
^1^H-NMR analysis of RG-II isolated from. **a**, Col-0 wildtype, **b**, NE control, **c**, *rckt1-9* and **d**, *rckt1-21*. The isolated RG-II (dimers) was treated with dilute HCl to generate monomers for the analysis. The diagnostic resonance of Kdo is shown in **a** and **b**, but absent from **c** and **d**. The Dha signal is detected in all the samples.

**Fig 5.**
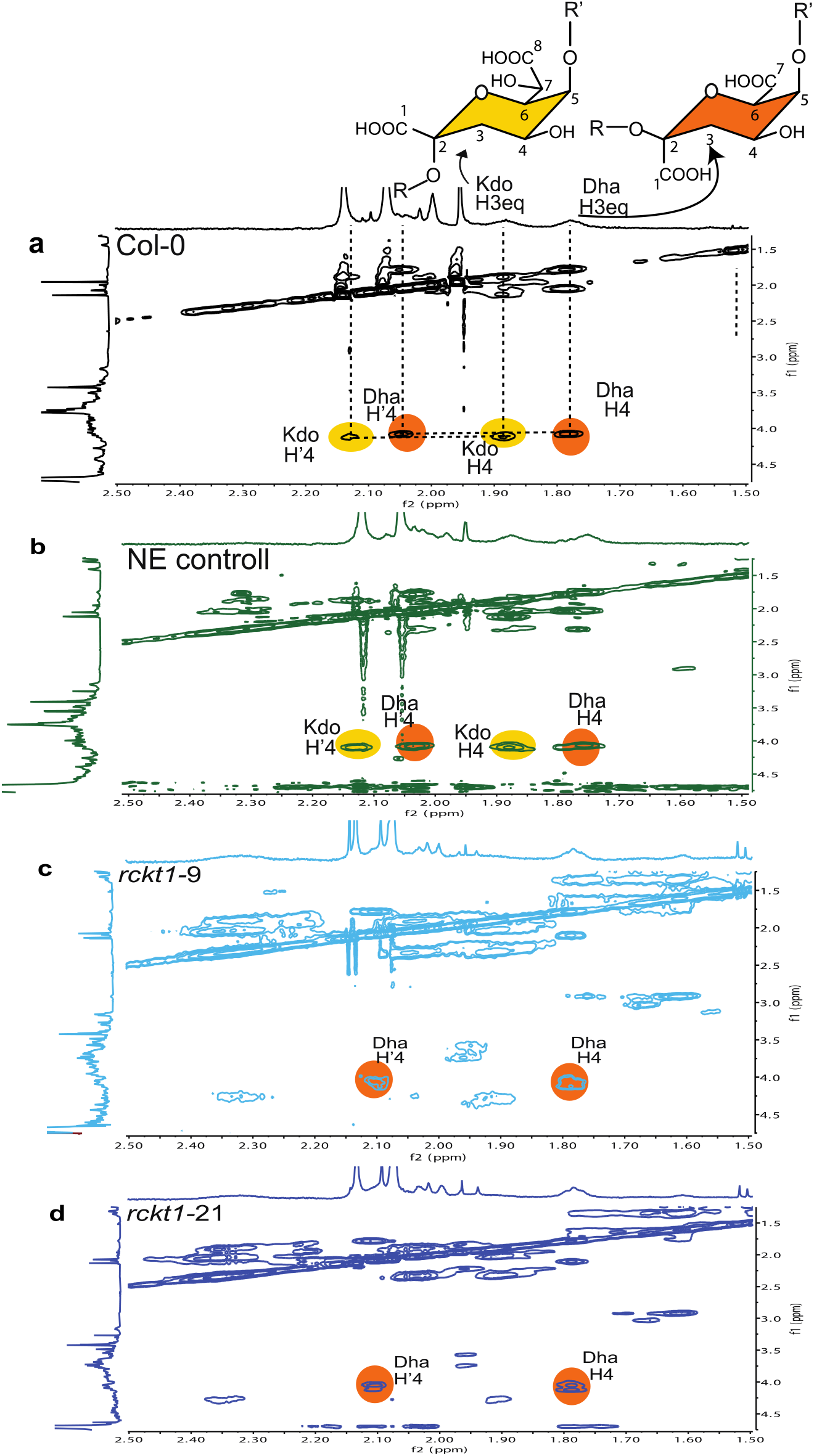
2D TOCSY NMR analysis of RG-II isolated from. **a**, Col-0 wildtype, **b**, NE control, **c**, *rckt1-9* and **d**, *rckt1-21*. The isolated RG-II (dimers) was treated with dilute HCl to generate monomers for the analysis. The diagnostic resonance of Kdo is shown in **a** and **b**, but absent from **c** and **d**. The Dha signal is detected in all the samples.

We next prepared the trimethylsilyl methyl-ester methyl glycoside (TMS) derivatives of the RG-IIs for analysis by gas-chromatography with electron impact mass spectrometry (GC-EI-MS) ^45^ . Dha is present in the controls and the *rckt1* mutants, whereas Kdo is detected in the controls, but absent in the *rckt1-21* homozygous mutant and the *rckt1-9* heterozygous/biallelic mutant (**Supplementary** Fig. 4). In summary, our NMR spectroscopic and glycosyl residue composition analyses conclusively demonstrate that the homozygous mutant and the *rckt1-9* heterozygous/biallelic mutant produce RG-II that lacks discernible amounts of Kdo. This evidence supports our hypothesis that RCKT1 is the Kdo transferase required to form RG-II side-chain C.

### Predicted structure of RCKT1 in the presence of CMP-β-Kdo

Determining the biochemical properties of RCKT1 is challenging since its acceptor is unknown and the CMP-β-Kdo donor has an estimated half-life of ∼34 minutes ^47, 48^. Thus, we used AlphaFold ^49^ to predict the conformation RCKT1 and gain some molecular insight into its function. The model shows a N-terminal transmembrane domain from residue 1 to 36 with the transmembrane ɑ-helix between residues 12 and 36. RCKT1 is a mix of ɑβ folds, comprising 1 central β-sheet consisting of 7 parallel β-strands surrounded by 14 ɑ-helices and 2 small β-sheets (**Fig. 6a-b**). The mammalian α-2,3-sialyltransferase ST3Gal I (PDB ID 2WNB) from *Sus Scrofa* (wild pig) ^50^ shares 32% amino acid sequence identity with RCKT1 and represents the closest structural homolog in the Protein Data Bank. Both enzymes are members of the GT29 family, whose structurally characterized members have been shown to adopt a modified GT-A fold, but lack the characteristic DxD motif ^51^. The structure of the GT42 sialyltransferase CstII (PDB ID 1RO7) from *Campylobacter jejuni* ^52^, which has similar folding to RCKT1, was solved in complex with CMP-3FNeuAc. A superposition of CStII and RCKT1, allowed us to identify the likely position of the CMP binding site and consequently model CMP-β-Kdo into the substrate donor active site of RCKT1. This putative binding site suggests that the main and side chain atoms of Phe306, Thr307 and Thr316 directly contact the cytidine nucleotide. Moreover, Trp315 makes aromatic stacking interactions. The ribose portion of the substrate interacts with the main chain N atoms of Asn186 and Gly287, and the Kdo sugar region showed 2 hydrogen bonds with Thr236 (**Fig. 6c**). It will be important to test the importance of these predicted residues in future experiments.

**Fig. 6.**
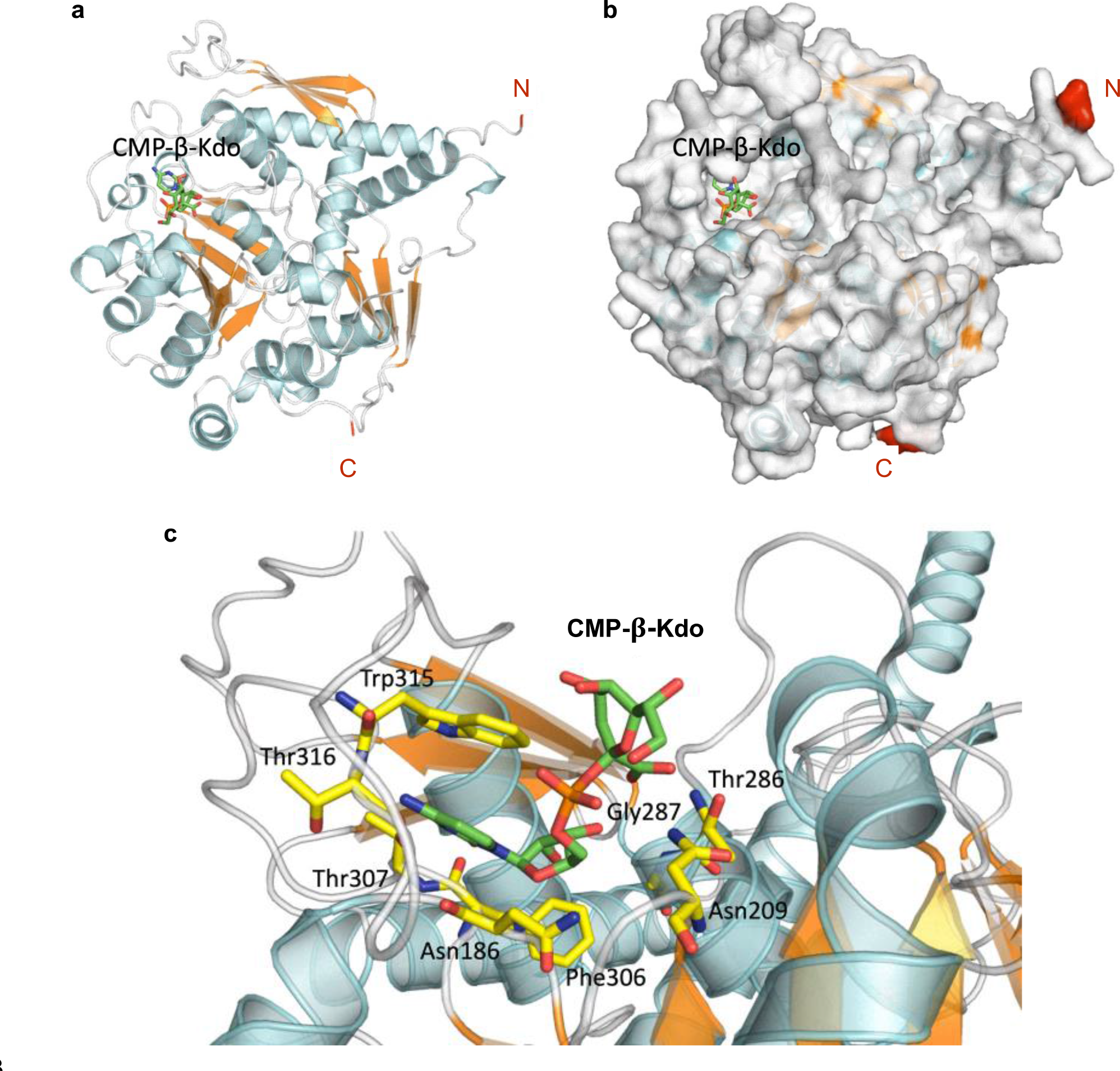
Structure of Arabidopsis RCKT1 predicted by AlphaFold modeling. **a**, Cartoon representation of general architecture of RCKT1 model. The predicted N-terminal transmembrane domain correspondent from residue 1 to 36 is not shown. The nucleotide donor substrate, CMP-β-Kdo, is positioned atop a central β-sheet comprising seven parallel β-strands surrounded by 14 ɑ-helices and 2 small β-sheets. **b,** Surface representation of RCKT1 structure in complex with CMP-β-Kdo. It reveals the potential CMP-β-Kdo binding site and the surrounding binding cleft, which is a likely location for the binding of the acceptor substrate. **c**, Zoom in on the CMP-β-Kdo binding site showing all the residues interacting via hydrogen bond and aromatic interactions between RCKT1 and CMP-β-Kdo ligand.

## Discussion

### RG-II structure integrity is important for its function

It is widely recognized that the structure of RG-II must be maintained to ensure it can form a dimer normally and that this is a key factor in its functions related to plant growth, development, and viability. This view has primarily been supported by research on plant mutants deficient in RG-II specific GTs, enzymes and transporters responsible for RG-II specific nucleotide sugars, and boron uptake/transport mutants. Knockdown of the RGXTs involved in side-chain formation ^26^, and disruption of the synthesis and transport of RG-II nucleotide sugars, such as GDP-L-Fuc ^15–17^, GDP-L-Gal ^18, 19^, UDP-D-Api ^20^, all lead to severe growth and developmental defects in plants. The truncation of side-chain A, due to the absence of D-Xyl, L-Fuc, L-Gal or D-Api is believed to be responsible for the defective RG-II dimerization. These data are consistent with the notion that the borate cross-link formed between two RG-II monomers involves the chain A by D-Api residue^3, 4^.

Our study of RCKT1 introduces a new perspective by showing that preventing the formation of side-chain C, a disaccharide seemingly distant from the boron-binding D-Api residue, also disrupts dimer formation. This broadens the prevailing hypothesis that any alteration to the RG-II structure may have a substantial impact on its dimerization and compromise the formation of a normal pectic network *in muro*. Our CRISPR-mediated strategy provides a basis to test this hypothesis. Generating loss-of-function mutants of RG-II GTs, would allow us to create a diverse array of RG-II glycoforms *in situ*, allowing for an exploration of the relationship between RG-II structure and functionality. Beyond RG-II dimerization, we can also delve into whether structural modifications of RG-II influence potential interaction with other pectic domains and various cell wall components. This would open new avenues for us to probe how the pectic network contributes to the mechanical strength and physical properties of the plant primary wall, which play a substantial role in shaping the growth and development of plant cells, tissues, and organs ^53–55^. The formation of a pectic network involves multiple levels of crosslinking, including but not limited to the backbone glycosidic linkages of the three pectic domains, the borate ester crosslinking of RG-II, and the calcium crosslinking of HG ^56^. Although the specific role of RGII- boron cross-linking in maintaining the physical property of plant cell wall is far from clear, it has been suggested to regulate wall pore size ^57^, mechanical strength ^58^ and extensibility ^1^.

### Challenges to confirm the enzymatic activity of RCKT1 *in vitro*

Our data have shed light on the role of RCKT1 in the addition of Kdo to RG-II. An *in vitro* assay for enzymatic activity will be required to increase our understanding of RCKT1’s function as an RGII-specific Kdo transferase. However, this is challenging for several reasons including the difficulties of heterologously expressing a functional RCKT1 enzyme, the availability of the donor substrate CMP-β-Kdo, and the availability of appropriate RG-II glycoforms as acceptor substrates.

Plant cell wall GTs have been challenging to express as stable and active recombinant proteins in bacteria and yeast ^59, 60^. Our attempts to express RCKT1 using the *Escherichia coli* Origami B strain was not successful. The Human Embryonic Kidney (HEK293) cell system ^61^, provides an alternative technology as it has been used to produce several plant GTs ^62–68^. However, an equally large challenge arises from the instability of CMP-β-Kdo, the donor substrate for the RCKT1 enzyme. CMP-β-Kdo is rapidly hydrolyzed in aqueous buffers (half-life of ∼34 mins) resulting in the formation of CMP and free Kdo ^47, 48^. To address this, several groups have generated CMP-β-Kdo *in situ* using CTP, Kdo and a CMP-β-Kdo synthetase (KdsB) purified from *E. coli*. This strategy has proven successful in studies focused on β-Kdo-transferases involved in bacterial glycolipid biosynthesis ^69–71^. Finally, an appropriate RG-II glycoform, capable of accepting Kdo from the nucleotide sugar donor CMP-β-Kdo, is required. It is possible that an RG-II lacking side-chain C could serve as an ideal acceptor for Kdo. The *RCKT1* callus mutant generated in this study could provide this glycoform. Alternatively, sidechain C could be removed using the bacterial glycanase described by Ndeh et al ^14^. However, the possibility cannot be discounted that a glycoform lacking other side-chains in addition to side-chain C is the acceptor. Tools are now available to modify RG-II structure *in vitro* and generate different glycoforms in amounts suitable for GT assays ^14^. Nevertheless, identifying the appropriate acceptor molecule remains a considerable challenge.

It will also be important to obtain structural information to understand the molecular mechanism of its catalytic activity. This will allow comparison of the sialytransferase motifs from the closely related animal STs with the plant RCKT1 to identify potential residues responsible for substrate specificity. Future work will include generating a series of RCKT1 variants for kinetic analysis. It will also be interesting to compare this protein to the previously identified AtKDTA KDO transferase candidate, a mitochondria-localized protein which is thought to have a role in glycosylating an as-yet unidentified lipid signaling molecule ^72^.

### Advantages of, and potential extensions to, a callus-based gene editing method

We have demonstrated the utility of our callus-based gene editing method for generating KO mutations in two GTs that are required for normal plant growth and development. We believe that this approach could serve as a tool for both reverse and forward genetic investigations. One application would be the creation of callus carrying mutations in multiple different GTs. Gene redundancy significantly hampers the characterization of plant GTs ^59^. For example, the GAUT family of Arabidopsis has 15 members. Only a few of these have yielded mutants with observable phenotypes that assist in revealing gene function ^27, 73, 74^. The multiplexed CRISPR/Cas9 toolbox employed in our study allows the assembly of up to eight sgRNAs into a single T-DNA vector, a crucial feature that can enable the simultaneous targeted editing of multiple genes ^42^.

Our method can be adapted for use in forward screens to identify candidate GTs involved in specific biological processes, such as cell-cell adhesion. Known cell adhesion mutants exhibit a decreased regeneration phenotype ^75^. It should be possible to create a library of CRISPR callus mutants for the 75 candidate genes in our target clade, and assess organ regeneration capability. The CRISPR/Cas9 system has been used with rice to generate a large-scale mutant collection ^76^.

Improvements to gene editing efficiency would enable this method to be more widely implemented. Only one of the three bioinformatically predicted top-performing sgRNAs created a CRISPR mutation at the target sites of *GMT1* and *RCKT1*. Liang and colleagues have developed a prescreening method to assess sgRNA efficiency via the tobacco (*Nicotiana benthamiana*) transient expression system ^77^, that could be applied to select highly efficient sgRNAs for the target genes. This is a rapidly developing area of research, with recent improvements including the use of stronger promoters to drive higher expression of the sgRNAs and Cas9, the utilization of intronized Cas9 ^78^, or alternative Cas enzymes, can be used to further optimize its editing efficiency in plant species.

## Materials and Methods

### Plant material and incubation conditions

*Arabidopsis thaliana* ecotype Col-0 and *gmt1-3* seeds were surface sterilized for 10 minutes using a 10% (v/v) commercial bleach (Clorox, 6.5%, v/v) in aq. Ethanol (100% v/v), and then washed twice with 100% ethanol, and allowed to air dry overnight. Sterilized seeds were placed on B5 agar media (**Supplementary Table 3**) and stratified for three days at 4°C in the absence of light. The plates were then transferred to a growth chamber (8,000 lux, 16:8 L:D, 22°C, 75% humidity). After 10 days, the primary roots were isolated from the seedlings for root transformation as described below. *gmt1-3* allele is as described in ^30^. The callus culture of *gmt1-3* was derived from roots as described in ^79^.

Calli were maintained by sub-culturing every 14 days onto freshly prepared callus induction media (CIM) (**Supplementary Table 3**). The callus selected for subculture should be creamy and white, indicating healthy status. Calli plates were kept in the dark in a growth chamber (25°C, 50% humidity). To grow cell suspension culture in liquid media, callus tissue (approx. 100 mg) was inoculated into 1 L flask containing 250 mL liquid CIM media and shaken at 100 rpm in the dark at 23°C, a method modified from ^80^. Callus from liquid cultures was harvested after 14 days of growth. The tissue was collected into falcon tubes and lyophilized at −50°C for 5 days. Fully lyophilized callus tissue was stored at −80°C for downstream biochemical analysis.

### Screening GTs important for plant cell-cell adhesion by a bioinformatic pipeline

To identify putative GTs involved in the plant cell-cell adhesion process, we set up a bioinformatics pipeline based on the following filtering process.

Filter I: Arabidopsis GT genes were obtained from the CAZy database (http://www.cazy.org/). Genes corresponding to family GT1 proteins were removed since they are largely involved in the glycosylation of small molecules and are unlikely to be involved in cell wall biosynthesis. 441 genes from 49 GT families were identified.

Filter II: At*GMT1*, which encodes a GIPC mannosyltransferase was selected as the query gene for co-expression analysis. Co-expression analysis used 113 developmental RNA seq datasets across 11 tissue and organ types of Arabidopsis, and ThaleMine (https://bar.utoronto.ca/thalemine/begin.do) ^81^. Genes with similar expression patterns across the datasets were grouped based on hierarchical clustering using Cluster 3.0 (http://bonsai.hgc.jp/~mdehoon/software/cluster/). The gene clusters were visualized using TreeView (https://jtreeview.sourceforge.net/) and the clade containing the query gene *GMT1* was extracted.

Filter III: Genes encoding GTs localized in Golgi. The subcellular localization of the extracted GTs was obtained by SUBA (https://suba.live/). This step resulted in 75 Golgi-localized candidates as putative GTs most likely to be relevant to plant cell-cell adhesion.

### Guide RNA design and transformation vector construction

For the three sgRNA designed to target each gene, DNA oligonucleotides were synthesized by Integrated DNA technologies (**Supplementary Table 4**) and then annealed and phosphorylated. The phosphorylated sgDNA were individually cloned into the BsmBI site of the Golden Gate entry vectors pYXA or pYXB to create pY1A-AtU6-gRNA1, pYPQ132B-AtU3-gRNA2, and pY3A-AtU6-gRNA3. The three gRNA expression cassettes were then cloned into the recipient vector pYPQ143Amp by Golden Gate assembly to create a vector that contains the Tri- sgRNA module. The Tri-sgRNA module and Cas9 were assembled into the pTkan destination vector by Gateway cloning.

This modular cloning toolkit for CRISPR/Cas9 vector construction was originally developed by Lowder et al. ^42^, and modified in this study. The antibiotic selection markers of the Golden Gate recipient vector pYPQ143 and the Cas9 entry vector pYPQ150 from the original toolkit were replaced by the ampicillin resistance marker (AmpR) to be compatible with the pTKan destination vector we had in house.

Whole plasmid sequencing was performed to validate the sequences of the different components in the transformation vectors targeting AtGM1 and AtRCKT1. The final constructs were transformed into *Agrobacterium fabrum* (previously known as *Agrobacterium tumefaciens*) GV3101 for Arabidopsis root transformation.

### Arabidopsis root transformation

*Arabidopsis thaliana* ecotype Col-0 roots were transformed with *A. fabrum* GV3101 carrying the CRISPR/Cas9 vectors. The method was modified from Vergunst et al. ^41^. Sterilized Col-0 seeds were sown on B5 agar media (**Supplementary Table 3**) and grown in a growth chamber (8,000 lux, 16:8 L:D, 22C°, 75% humidity). Primary roots were isolated from 10d old seedlings using a sterile razor blade and transferred onto CIM media (**Supplementary Table 3**) and grown for three days. *A. fabrum* GV3101 carrying the transformation vector was grown for two days at 30°C on LB solid media with antibiotics (rifampin, gentamicin, spectinomycin). A single *A. fabrum* colony was inoculated into liquid LB media with antibiotics (rifampin, gentamicin, spectinomycin) and cultured overnight at 30°C with shaking at 200 rpm. *A. fabrum* cells were pelleted and then suspended in liquid B5 media (final OD600 = 0.1). The isolated primary roots were soaked for five minutes in the *A. fabrum* suspension, then sliced into 0.5-1 cm explants using a sterile razor blade. The root explants were blot dried on sterile filter paper and plated onto CIM media containing D-glucose (1.8 g/L) and acetosynringone (0.1mM). Root explants were co-cultivated with *A. fabrum* for three days at 25°C under dim light (2,000 lux, 16:8 L:D). The root explants were then washed four times with Timentin-supplemented (100mg/L) B5 liquid media and plated onto CIM media supplemented with antibiotics (50 mg/L Kanamycin and 100mg/L Timentin). After 14 days, the explants were transferred to fresh CIM media containing Kanamycin and Timentin. Calli resistant to Kanamycin were collected and sub-cultured every 14 days.

### CRISPR editing analysis of the transgenic callus mutants

Callus tissue was sampled from each independent transformation event, flash frozen in liquid nitrogen, and then milled with metallic beads using a TissueLyzer (Qiagen) for subsequent genomic DNA (gDNA) extraction. Genomic DNA was extracted from callus using the QIAGEN DNeasy Plant Mini Kit®. Primers that amplified the genomic region of the guide RNA (gRNA) target sites were used to genotype the material (**Supplementary Table 5**). PCR amplicons were column-purified using the Zymo Research DNA Clean & Concentrator® kit, and Sanger sequenced by Azenta (https://www.azenta.com/). Decodr v3.0 (https://decodr.org/) was used to deconvolute the sequenced amplicon for mutation analysis.

### GIPC purification and analysis by TLC

GIPC analysis by TLC was performed as described by Jing et al. ^44^. Lyophilized callus tissue (∼0.5 g dry weight) harvested from three distinct liquid cultures were combined and ground to a powder with metallic beads using a TissueLyzer (Qiagen). Sphingolipids were extracted from the ground tissue by using the lower layer of the isopropanol/hexane/water (55:20:25 v/v/v) extract followed by deesterification with 4 mL of 33% methylamine solution in ethanol/water (7:3 v/v) as previously described ^82^. The water used in this experiment is HPLC-grade. To enrich the GIPCs, the dried extracts were suspended in 1 mL of enrichment solution (chloroform/ethanol/ammonia/water [10:60:6:24 v/v/v/v]) and kept overnight at room temperature with agitation (2 rpm). The GIPCs were then isolated using a weak anion exchange SPE cartridge (Strata X-AW; Phenomenex) as described ^29^. GIPCs bound to the cartridge were eluted with 3 mL of enrichment solution and dried under a flow of nitrogen gas or a stream of clean dry air. The dried GIPCs were suspended in solution A {chloroform/methanol/[4M ammonium hydroxide in 1.8M ammonium acetate] (9:7:2 v/v/v)} and separated by thin layer chromatography (TLC). The separations were performed on high-performance TLC Silica gels on glass plates (Merck) with solvent A as the mobile phase. GIPCs were stained with primuline (10 mg in 100 mL acetone/water [8:2 v/v]), and visualized under a fluorescent blue light (460 nm) ^83^.

### GIPC glycan profiling by HPAEC-PAD

Purified GIPCs were treated for 1 hour at 120°C with 2M trifluoroacetic acid (TFA, 1 mL), cooled on ice, and dried overnight using a vacuum concentrator. The residues were dissolved (2-5 fold) in HPLC grade water and the liberated monosaccharides analyzed by high-performance anion-exchange chromatography with a pulsed amperometric detector (HPAEC-PAD) as described ^30^.

### Preparation of plant cell walls as alcohol insoluble residue (AIR)

AIR was prepared as previously described ^29^. Briefly, lyophilized callus was milled into a fine powder and then suspended in 96% (v/v) ethanol. The suspension was kept for 30 min at 70°C, and the suspension then cooled and centrifuged (15 mins, 18,500 rcf). The pellet was then suspended in ethanol (96% v/v), vortexed, and then centrifuged again (15 mins, 18,500 rcf). The pellet was then suspended in methanol:chloroform (2:3 v/v) and kept overnight at 4 °C with agitation (300 rpm). Following centrifugation (15 min, 18,500 rcf), the pellet was treated for 1h at 4°C methanol:chloroform (2v:3v) (1h, 300 rpm,). Following centrifugation (15 min, 18,500 rcf), the pellet was washed with ethanol: 100% (v/v), 65% (v/v), 70% (v/v), 80% (v/v), 100% (v/v). Each wash involved a 5-minute centrifugation (18,500 rcf) and removal of the supernatant in between, with a thorough vortex to suspend the pellet. The resulting alcohol insoluble residue (AIR) was dried at 50 °C overnight in a vacuum concentrator or oven.

Starch was removed by treating the AIR (∼10 mg) for 10 mins at 75°C with agitation at 300 rpm) with *α*-amylase (3 U/mL) in 50 mM MOPS, pH 7.3). Pullulanase (1 U/mL) and amyloglucosidase (3 U/mL) X mM Na acetate, pH 4.5, were then added, and the suspensions kept for a further two hours at 40°C (with agitation at 300 rpm). Following centrifugation, the pellet was washed with the following ethanol series: 100% (v/v), 70% (v/v), 70% (v/v). The destarched AIR was dried overnight at 30°C in a vacuum concentrator.

### Cell wall monosaccharide composition analysis

Non-cellulosic sugars were liberated from de-starched AIR by TFA hydrolysis for one hour at 120°C. The supernatant containing the liberated monosaccharides was collected after centrifugation. Residual monosaccharides in the remaining pellet were further extracted using water and combined with the supernatant. The combined supernatant was dried overnight at 30°C in a vacuum concentrator. The residue was suspended in HPLC-grade water and filtered through a nitrocellulose filter (0.45 μM). The non-cellulosic sugars in the filtrate were diluted (4-5 fold) with water and filtered through a nitrocellulose filter (0.45 μM) prior to HPAEC-PAD analysis ^30^.

The TFA-insoluble AIR pellet was then subjected to Saeman hydrolysis to hydrolyze cellulose. The pellet was suspended in 72% (v/v) sulfuric acid and kept for one hour at room temperature. Water was added to dilute the sulfuric acid to 1M and the suspensions then heated for 3h at 100°C. The suspensions were then neutralized by adding barium carbonate and centrifuged. The supernatant containing was dried overnight at 30°C in a sample concentrator. The residue was suspended in water and filtered through a nitrocellulose filter (0.45 μM). The sugars in the filtrate were diluted with water and filtered prior to HPAEC-PAD analysis. The water used in this experiment is HPLC-grade only.

### Isolation of RG-II from AIR of controls and the *rckt1* mutants

The callus AIR (500-700mg) extracted from three distinct liquid cultures were combined and suspended in NaOAc (50 mM, pH 5.0) at a concentration of 25mg/ml, and then treated for 24 hrs at 37 °C with 5 U/g endopolygalacturonase (EPG) and 10 U/g of pectin methyl esterase (PME) with shaking at 150 rpm. The EPG solublized material and the insoluble residue were separated by filtration through a nylon mesh (100 μm). The insoluble material retained on the mesh were treated again with EPG and then filtered as before. The EPG-soluble materials were obtained by combining the filtrates after the two treatments. The x mL solution in total were then fractionated by preparative size-exclusion chromatography (SEC) on a column Superdex Increase 75, (10/300 GL, Cytiva, USA, column L × I.D. 30 cm × 10 mm, 9 μm) with refractive index (RI) detection. The column was eluted with ammonium formate (50 mM, pH 5.0) at 0.5 mL/min. Fractions enriched in RG-II were collected manually, dialyzed (3500 Dalton MWCO) against deionized water, and freeze-dried.

### RG-II glycosyl residue composition analyses

To specifically detect Kdo and Dha, the glycosyl residue of RG-II were analyzed as their trimethylsilyl (TMS) methyl glycoside derivatives by gas-chromatography with electron impact mass spectrometry (GC-EI-MS) as described in ^45^. In general, RG-II (100 μg) was suspended in 300 μL methanolic HCl (1 M) in screw-top glass tubes secured with Teflon-lined caps and heated at 80 °C for 18 hrs. After cooling to room temperature, the solutions were concentrated to dryness under the nitrogen stream. The released methyl glycosides and methyl glycoside methyl-esters were then reacted for 30 min at 80 °C with Tri-Sil^®^ (ThermoFisher, USA). GC-EI-MS analysis of the TMS methyl glycosides was performed on an Agilent 7890A GC interfaced to an Agilent 5975C mass selective detector, with a Supelco Equity-1 fused silica capillary column (30 m × 0.25 mm ID).

### Preparation of RG-II monomer for NMR spectroscopic analysis

The RG-II dimer was converted to the monomer as previously described ^45^. Briefly, the RG-II enriched material obtained by SEC was treated for 1 h at room temperature with HCl (0.1 M). The solution was then dialyzed (3500 Dalton MWCO) against deionized water and freeze-dried.

### 1D ^1^H-NMR and 2D TOCSY spectroscopy of RG-II monomer

RG-II (1-2 mg) was dissolved in D2O (0.2 mL, 99.9%; Cambridge Isotope Laboratories, Tewksbury, MA, USA) and placed in a 3 mm NMR tube ^45^. Spectra were obtained using a Bruker NMR spectrometer (Agilent Technologies) operating at 900 MHz using a 5 mm cold probe and a sample temperature of 25°C. The 1D ^1^H and 2D TOCSY spectra recorded using standard Bruker pulse programs. Chemical shifts of the RG-II glycosyl residue were measured relative to internal D2O (δH 4.707). Data were processed using MestReNova software (Mestrelab Research S.L., Santiago de Compostela, Spain).

## Statistical analysis

One-way ANOVA were used for statistical analysis.

## Accession Numbers

Sequence data from this article can be found at The Arabidopsis Information Resource (TAIR) under accession number At3g55830 (*AtGMT1*), At1g08660 (*AtRCKT1*), AT3G48820 (*AtSIA2*), and AT1G08280 (*AtGALT29A*).

## Acknowledgements

This work was part of the DOE Joint BioEnergy Institute (http://www.jbei.org) supported by the 1. U. S. Department of Energy, Office of Science, Office of Biological and Environmental Research, through contract DE-AC02-05CH11231 between Lawrence Berkeley National Laboratory and the U. S. Department of Energy. This work was also supported in part by an establishment grant to JCM from the University of Adelaide. Work conducted at the Complex Carbohydrate Research Center, of the University of Georgia, was supported by the Division of Chemical Sciences, Geosciences, and Biosciences, Office of Basic Energy Sciences of the United States Department of Energy through Grant DESC0008472 for funding structural studies of RG-II and DE-SC0015662 to the Complex Carbohydrate Research Center. We thank Maartje van Kregten and Annette Vergunst for the insightful discussion on the Agrobacterium-mediated root transformation of Arabidopsis. We thank Yu Gao and Yi-Chun Chen for their support with HPAEC-PAD experiments. We thank Jutta Dalton and Mi Yeon Lee for their support with growth chamber maintenance. We thank Ramana Pidatala for his advice on plant tissue culture. We thank Ariel Orellana for the insightful discussion on the putative CMP-Kdo transporter, homologous to CMP-sialic acid transporter identified in mammals. We thank Parastoo Azadi and the analytical service laboratory at the Complex Carbohydrate Research Center for technical support.

## Contributions

Y.Z. and J.C.M. conceived and designed the study. Y.Z. conducted the gene editing, transformation and growth experiments, assisted by N.D., Y.L., and F.D. YZ conducted GIPC and cell wall analysis. D.S., K.T., M.O., I.B., and B.U. structurally characterized RG-II. J.H.P. and P.A. performed AlphaFold modeling of RCKT1. Data analysis and interpretation were conducted by Y.Z., H.V.S. D.S., B.U., M.O., J.H.P., and J.C.M. The paper was written by Y.Z. and J.C.M. with revisions from all authors.

## Ethics declarations

The authors declare no competing interests.

## Data availability

The data supporting the findings of this study are available within the paper and its supplementary information files.

