## Supplemental Information for "A putative rhamnogalacturonan-II CMP-β-Kdo transferase identified using CRISPR/Cas9 gene edited callus to circumvent embryo lethality"

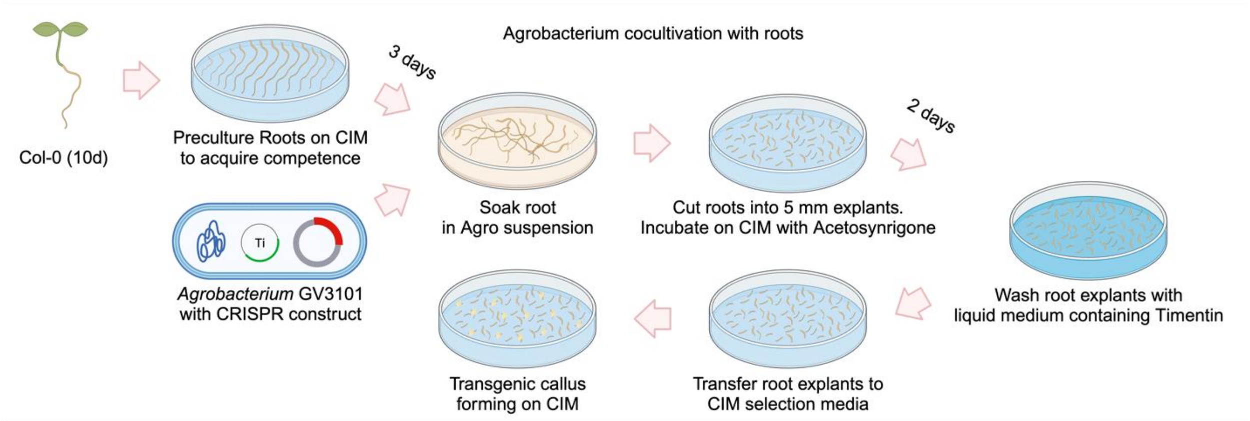


**Supplementary Fig. 1| *Agrobacterium*-mediated Root Transformation of Arabidopsis.** Schematic illustration of the transformation procedure of Arabidopsis root system to generate transgenic callus. Primary roots are taken from 10-day-old Arabidopsis seedlings and pre-cultured on callus induction media (CIM). After three days, the primary roots are infected with Agrobacteria carrying the T-DNA vector and then cut into root explants. The infected root explants are incubated on co-cultivation media for two days, and then washed using liquid media supplemented with Timentin (100 mg/L) to neutralize the infecting Agrobacteria. The explants are then incubated on CIM media supplemented with selection antibiotics (Kanamycin) only allowing transgenic callus with stable T-DNA integration to survive after about one month.

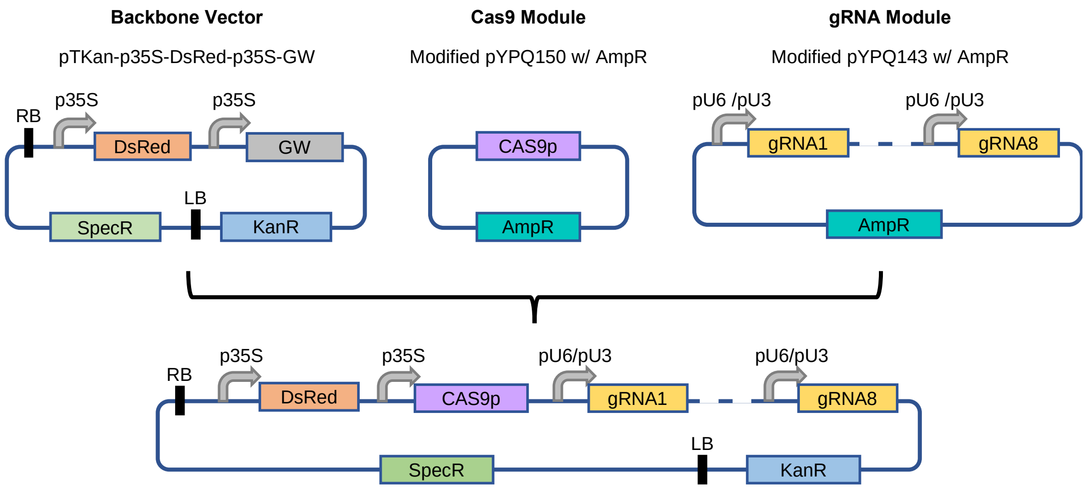


**Supplementary Fig. 2| Tailored CRISPR/Cas9 platform for GT editing in Arabidopsis.** The assembly procedure of CRISPR/Cas9 T-DNA vector is adapted from (Lowder et al. 2015)**.** Three modules are required for this platform: a T-DNA destination vector providing 35S promoter for Cas9 expression; a Cas9 entry vector; a gRNA entry vector that contains up to eight gRNA expression cassettes. The T-DNA transformation vector can be made by the Multisite Gateway recombination to assemble the Ca9 and the gRNA modules into the T-DNA destination vector. The module containing multiple gRNA expression cassettes is made by a two-step Golden Gate cloning as described in [42]. SpecR, spectinomycin resistance marker; KanR, kanamycin resistance marker; DsRed, DsRed fluorescence reporter; GW, Gateway recombination sites; 35S, CaMV 35S promoter; LB, left border region; RB, right border region; AmpR, ampicillin resistance marker; pU6, Arabidopsis ubiquitin6 promoter; pU3, Arabidopsis ubiquitin3 promoter.

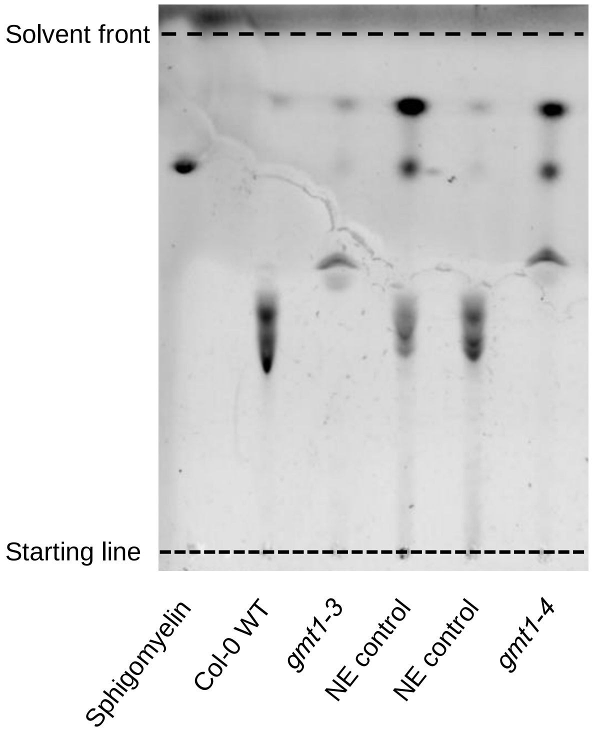


**Supplementary Fig. 3| GIPC analysis using TLC.** GIPCs were extracted from Arabidopsis callus tissue of Col-0 wild-type, *gmt1-3*, NE control, and *gmt1-4*. The enriched GIPCs and sphingomyelin standard were run on a TLC plate and stained with Primuline before imaging.

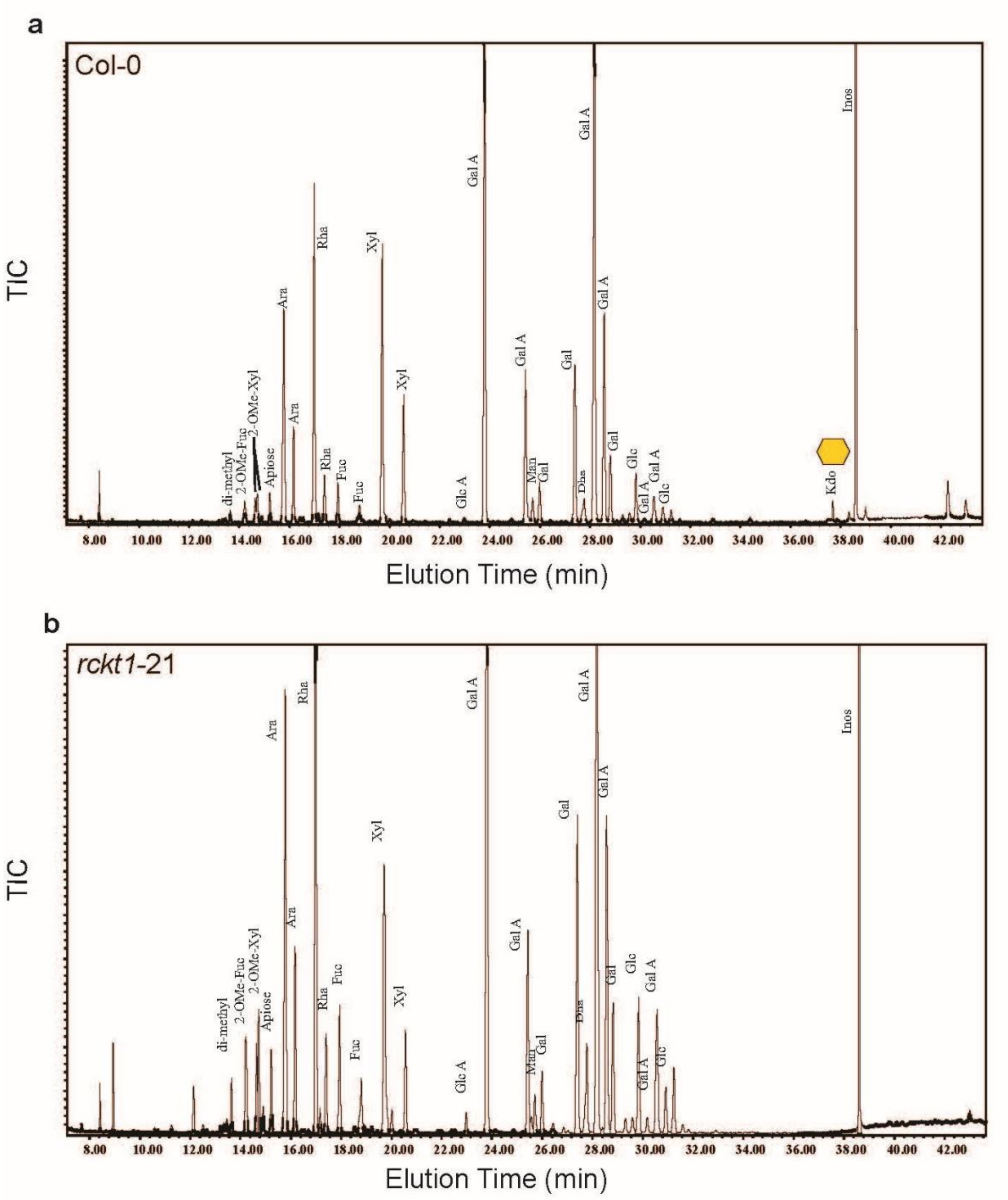


**Supplementary Fig. 4| GC-EI-MS total ion current profiles of the trimethylsilyl methyl-ester methyl glycoside derivatives of the monosaccharides generated from RG-II.** The GC-EI-MS total ion current (TIC) profile of the TMS derivatives generated from RG-II of **a**, Col-0 wild-type, **b**, *rckt1-21*. The identity of the monosaccharide derivative in each peak is shown. The peak eluting at ~ 39 min is the TMS derivative of Myo-inositol used as an internal standard.

**Supplementary Table 1**| Single guide RNA (sgRNA) sequences for the target genes.

| **Target gene** | **gRNA sequence (5’ -> 3’)** | |
|---|---|---|
| ***GMT1*** | gRNA1 | TACGCAAGTTCGTGACGGCG |
| | gRNA2 | GTTCAATCGCCGTCGCAGAT |
| | gRNA3 | GCATGAGGTTGAGCTGAGAT |
| ***RCKT1*** | gRNA1 | GTCCAGCGTTCAACAGTGCG |
| | gRNA2 | CTGAAGGCACTAACAGTACT |
| | gRNA3 | TGTACGCCCTGATGGGTGGT |

**Supplementary Table 2**| **Mannose content of the GIPCs isolated from Arabidopsis callus tissue.** The GIPCs were extracted from a pooled collection of callus tissue obtained from three independently grown liquid culture for each line. Following extraction, the enriched GIPC fraction was hydrolyzed by TFA to liberate monosaccharides within their glycan headgroups. The derived monosaccharides were quantified by HPAEC-PAD. The mannose quantity was normalized to the dry weight (DW) of callus tissue.

| **Mannose content** | Col-0 | *gmt1-3* | NE control | *gmt1-4* |
|---|---|---|---|---|
| (mg/g DW) | 7 | 1 | 6 | 2 |

**Supplementary Table 3**| The abundance of the RG-II dimer and monomer in the material released by endopolygalacturonase (EPG) and pectin methyl esterase (PME) treatment of the alcohol-insoluble residue (AIR).

| Callus line | Released by EPG and PME treatment of AIR | |
|---|---|---|
| | Dimer | Monomer |
| | % of total RG-II[a] | |
| Col-0 | 70 | 30 |
| NE Control | 80 | 20 |
| *rckt1-9* | 26 | 74 |
| *rckt1-21* | 22 | 78 |

[a]The abundance of the dimer and monomer was determined by size-exclusion chromatography (SEC) with RI detection.

**Supplementary Table 4**| Composition of *Arabidopsis thaliana* tissue culture media.

| Components | | B5 | Callus induction media (CIM)[a] |
|---|---|---|---|
| Gamborg B-5 Basal Medium | | 3.2 g/L | 3.2 g/L |
| D-Glucose | | | 20 g/L |
| MES | | 0.5 g/L | 0.5 g/L |
| Difco Agar, Bacteriological | | 9 g/L | 9 g/L |
| Phytohormones | 2,4-D | | 0.5 mg/L |
| | Kinetin | | 0.05 mg/L |

[a]For co-cultivating Col-0 root explants with *A. fabrum*, CIM media with reduced glucose content (1.8 g/L) is supplemented with 0.1mM acetosynringone. For selecting transgenic callus line, CIM media is supplemented with 50 mg/L Kanamycin and 100mg/L Timentin.

**Supplementary Table 5.** Sanger sequencing primers for genotyping CRISPR callus mutants.

| **Primers** | **Sequence (5’ -> 3’)** |
|---|---|
| GMT1 Seq Primer_F | CACAGTTGACTTCTGAGACG |
| GMT1 Seq Primer_R | CAGAACGTGTGTGTGTGTAC |
| RCKT1 Seq Primer_F | GAGATCTCACCAAGCTGGCC |
| RCKT1 Seq Primer_R | GTCGAAGTGTCGGTAAACTC |
